# Automated Xeno-Free Chondrogenic Differentiation from Human Embryonic Stem Cells: Enhancing Efficiency and Ensuring High-Quality Mass Production

**DOI:** 10.1101/2024.05.20.594905

**Authors:** JunLong Chen, Oki Kataoka, Kazeto Tsuchiya, Yoshie Oishi, Ayumi Takao, Yen-Chih Huang, Hiroko Komura, Saeko Akiyama, Ren Itou, Masafumi Inui, Shin Enosawa, Hidenori Akutsu, Makoto Komura, Yasushi Fuchimoto, Akihiro Umezawa

**Author notes:** These two authors contributed equally to this work (JunLong Chen, Oki Kataoka).

## Abstract

**Introduction:** Repairing damaged cartilage poses significant challenges, particularly in cases of congenital cartilage defects such as microtia or congenital tracheal stenosis, or as a consequence of traumatic injury, as the regenerative potential of cartilage is inherently limited. Stem cell therapy and tissue engineering offer promising approaches to overcome these limitations in cartilage healing. However, the challenge lies in the size of cartilage-containing organs, which necessitates a large quantity of cells to fill the damaged areas. Therefore, pluripotent stem cells that can proliferate indefinitely are highly desirable as a cell source. This study aims to delineate the differentiation conditions for cartilage derived from human embryonic stem cells (ESCs) and to develop an automated cell culture system to facilitate mass production for therapeutic applications.

**Methods:** Cartilage cell sheets were derived from human ESCs (SEES2, clinical trial-compatible line) by forming embryoid bodies (EBs) with either conventional manual culture or a benchtop multi-pipetter and an automated medium exchange integrated cell incubator, using xeno-free media. Cell sheets were implanted into the subcutaneous tissue of immunodeficient NOG mice to obtain cartilage tissue. The properties of cartilage tissues were examined by histological staining and quantitative PCR analysis.

**Results:** We have optimized an efficient xeno-free system for cartilage production with the conventional culture method and successfully transitioned to an automated system. Differentiated cartilage was histologically uniform with cartilage-specific elasticity and strength. The cartilage tissues were stained by alcian blue, safranin O, and toluidine blue, and quantitative PCR showed an increase in differentiation markers such as ACAN, COL2A1, and Vimentin. Automation significantly enhanced the efficiency of human ESC-derived chondrocyte differentiation. The number of constituent cells within EBs and the seeding density of EBs were identified as key factors influencing chondrogenic differentiation efficiency. By automating the process of chondrogenic differentiation, we achieved scalable production of chondrocytes.

**Conclusions:** By integrating the differentiation protocol with an automated cell culture system, there is potential to produce cartilage of sufficient size for clinical applications in humans. The resulting cartilage tissue holds promise for clinical use in repairing organs such as the trachea, joints, ears, and nose.

## 1. Introduction

Cartilage is a non-vascular connective tissue that provides structural support throughout the body, supporting various body movements due to its diverse structure, which includes hyaline cartilage, elastic cartilage, and fibrocartilage. It is primarily comprised of chondrocytes embedded within an extracellular matrix (ECM) consisting of collagen fibers (such as type II, type IX, and type XI collagen molecules) and proteoglycans like aggrecan, link protein, and glycosaminoglycan[1]. Due to the absence of blood vessels, which are essential for cell proliferation and differentiation in situ[2,3], cartilage has limited self-repair capabilities. As a result, it often fails to regenerate completely after injury, leading to progressive localized damage and debilitating conditions like osteoarthritis, exacerbated by the lack of specific agents for cartilage repair.

Cell therapy and tissue engineering present promising approaches in regenerative medicine to address the limited healing capacity of cartilage[4–6]. These strategies involve utilizing chondrocytes, adult mesenchymal stromal cells (MSCs), and pluripotent stem cells. Primary chondrocytes obtained from the human body or cultured for short periods can regenerate cartilage, but fibroblasts induced during expansion culture hinder development of their properties and ability to reshape cartilage[7]. MSCs offer promise for regenerating and repairing cartilage lesions due to their differentiation potential and ease of collection with minimal invasiveness[8–11]. However, challenges such as reduced proliferative capacity and enhanced differentiation into fibrocartilage tissue have been observed with extended culture[5,7]. Embryonic stem cells (ESCs) and induced pluripotent stem cells (iPSCs) are viable sources of chondrocytes with unlimited proliferative capacity and self-renewal[7,12–14]. Transplanting chondrocytes derived from human pluripotent stem cells into defective articular cartilage has shown promise in forming cartilage without teratomas or tumors[13–15]. However, challenges remain in achieving fully differentiated chondrocytes to minimize the risk of teratoma formation[16,17].

Automating the procedure for generating substantial quantities of cartilage is also imperative to fulfill the substantial demand for clinical applications[18]. We employed a benchtop multi-pipetter and an automated medium exchange integrated cell incubator for EB preparation and cell sheet culture to achieve automation. This study highlights the success of mass-producing cartilage for human clinical applications due to the introduction of an automated culture system.

## 2. Materials and methods

### Culture of human ESCs

Human ESCs (SEES2) were cultured on dishes coated with iMatrix-511 silk (892021, nippi) in StemFit medium (RCAK02N, Ajinomoto).[19,20] The dishes were coated with 1.7 μg/ml iMatrix-511 silk at 37°C for 1 h. The medium was changed daily. ESCs were detached with a 1:1 mixture of 0.5 mM EDTA and TrypLE™ Select (A1217701, Thermo Fisher Scientific) for 5 min and passaged at 550 cells/cm^2^ every week.

### Formation of embryoid bodies

To generate embryoid bodies, ESCs were dissociated using a 1:1 mixture of 0.5 mM EDTA and TrypLE™ Select and plated on 96-well Clear Round Bottom Ultra-Low Attachment Microplates (7007, Corning) using benchtop multi-pipetter (EDR-384SR, BioTec). The cells were cultured in XF32 medium (81% Knockout DMEM (10829018, Thermo Fisher Scientific), 15% Knockout Serum Replacement XF CTS (12618013, Thermo Fisher Scientific), 2 mM GlutaMAX (35050061, Thermo Fisher Scientific), 0.1 mM MEM non-essential amino acids solution (11140050, Thermo Fisher Scientific), Penicillin-Streptomycin (15070063, Thermo Fisher Scientific), 50 μg/ml L-ascorbic acid 2-phosphate (A8960, Sigma-Aldrich), 10 ng/ml heregulin-1β (080-09001, Fujifilm Wako Pure Chemicals Co., Ltd.), 200 ng/ml recombinant human IGF-1 (85580C, Sigma-Aldrich), and 20 ng/ml human bFGF (PHG0021, Thermo Fisher Scientific)). The first day of the passage was performed with medium containing Rho-associated protein kinase inhibitor Y-27632 (036-24023, Fujifilm Wako Pure Chemicals Co., Ltd.). Embryoid bodies were collected by benchtop multi-pipetter at 4 days after the passage.

### Generation of cell sheets

To generate cell sheets, 672 embryoid bodies were plated on a 100 mm dish. The 100 mm dish was coated with 0.3 mg/ml NMP collagen PS (301-84621, nippi) at 37°C for 1 h. The embryoid bodies were cultured in XF32 medium at 37°C in 5% CO_2_. The medium was changed every 3 days using an automated medium exchange integrated cell incubator (CellKeeper (Model SCALE-120ME), RORZE Lifescience Inc.) in the automated cell culture system.

### Generation of cartilage tissue

Male immunodeficient NOG (NOD.Cg-PrkdcscidIl2rgtm1Sug/ShiJic) mice aged 6 weeks (Charles River Laboratories, Inc., Wilmington, MA) were used in this study. The mice were anesthetized by inhalation of 3% isoflurane (099-06571, Fujifilm Wako Pure Chemicals Co., Ltd.). Under sterile conditions, a cell sheet or cartilage harvested from cell sheets was transplanted between the subcutaneous scapulae of mice. The cartilage tissue was removed after mice were euthanized by cervical dislocation during inhalation anesthesia with 3% isoflurane. After removal, the cartilage tissue was fixed with 20% formalin.

### Preparation for histological studies

The cell sheets were coagulated in iPGell (PG20-1, GenoStaff) following the manufacturer’s instructions and fixed in 4% paraformaldehyde at 4°C overnight. Fixed samples were embedded in a paraffin block to prepare thin cell sections. Deparaffinization, dehydration, and permeabilization were performed using standard techniques. Hematoxylin-eosin (HE) staining was performed with Carrazzi’s hematoxylin solution (30022, Muto Chemicals) and eosin Y (32053, Muto Chemicals). Alcian blue staining was achieved with alcian blue solution pH2.5 (40852, Muto Chemicals). Nuclei were counterstained with kernechtrot solution (40872, Muto chemicals). Toluidine blue staining was achieved with toluidin blue solution pH7.0 (40981, Muto Chemicals). SafraninO staining was achieved with 1% safraninO stain solution (42162, Muto Chemicals) and 0.02% fast green water solution (15151, Muto Chemicals). Nuclei were counterstained with Weigert’s iron hematoxylin solution (40342 and 40352 Muto chemicals).

### Alcian blue staining in culture dishes

Samples in the culture dish were fixed in saccomanno solution (86542, Muto Chemicals) at room temperature for 3 hours. After washing dishes in dH_2_O two times, incubate with alcian blue solution pH2.5 (40852, Muto Chemicals) at room temperature for 45 min and decoloration three times with a 4:3 mixture of ethanol (057-00456, Fujifilm Wako Pure Chemicals Co., Ltd.) and acetic acid (017-00256, Fujifilm Wako Pure Chemicals Co., Ltd.) for 10 min. After washing dishes in dH_2_O three times, images were captured immediately using an all-in-one fluorescence microscope (Keyence, BZ-X710).

### qRT-PCR

Total RNA was prepared using ISOGEN (311–02501, Nippon Gene, Tokyo, Japan). The RNA was reverse transcribed to cDNA using Superscript III Reverse Transcriptase (18080–085, Invitrogen; Thermo Fisher Scientific, MA, USA) with ProFlex PCR System (Applied Biosystems, MA, USA). Quantitative RT-PCR was performed on QuantStudio 12 K Flex (Applied Biosystems, MA, USA) using a Platinum SYBR Green qPCR SuperMix-UDG (11733046, Invitrogen; Thermo Fisher Scientific, MA, USA). Expression levels were normalized with the reference gene, UBC. The primer sequences are shown in Table 1.

**Table 1.**
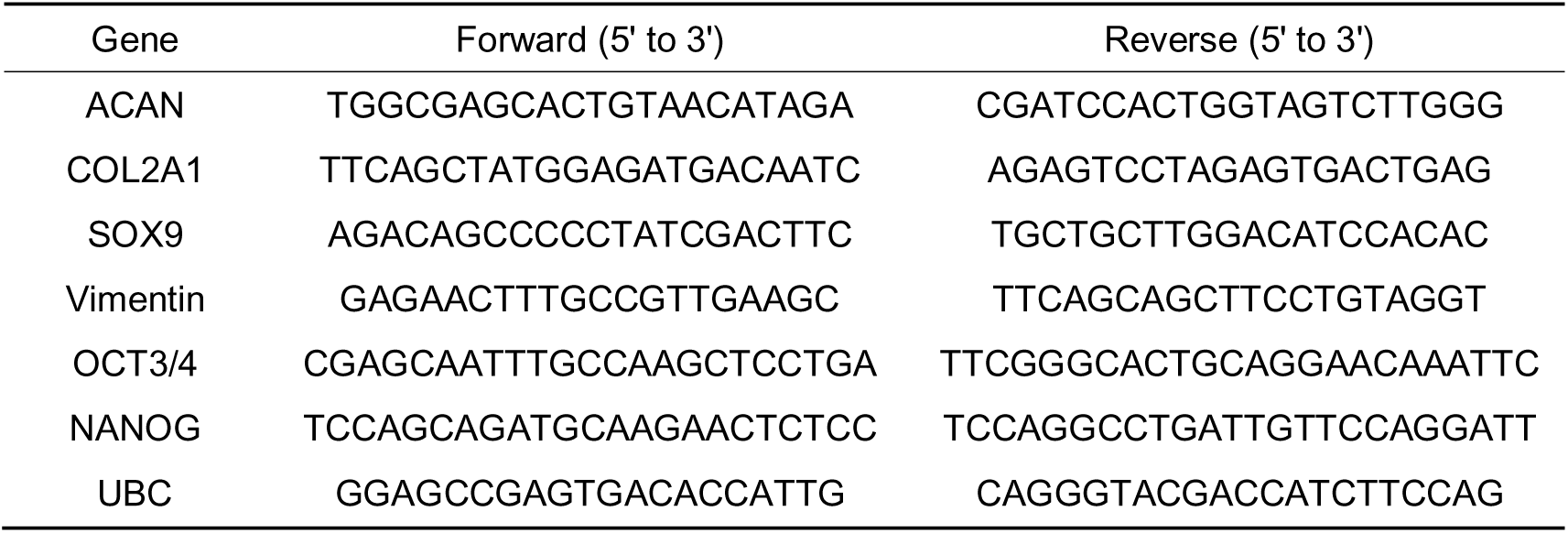

## 3. Results

### 3.1. A small number of EB-component cells facilitates chondrogenic differentiation

To obtain cartilage tissue, we generated cell sheets from hESCs and transplanted them into NOG mice (Fig. 1A). To improve the differentiation efficiency of the cell sheets, we optimized the culture conditions.

**Figure 1.**
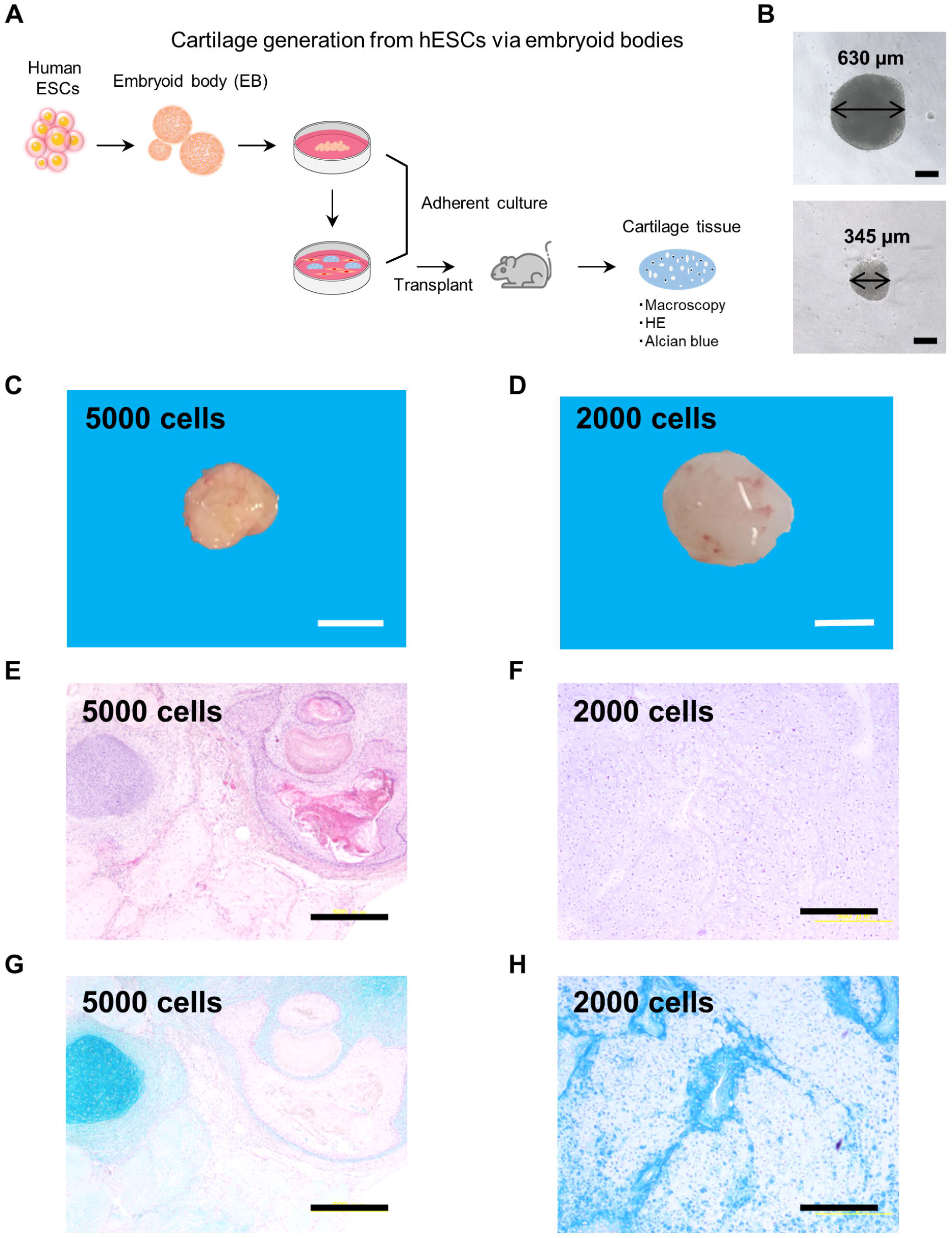
Effect of the number of EB constitutive cells on chondrogenic differentiation. A. Schematic diagram of the cartilage tissue differentiation protocol. B. Phase contrast microscopy images of EBs. Top: EB composed of 5000 cells, bottom: EB composed of 2000 cells; Scale bars = 200 µm. C. Macroscopic views of cartilage tissue differentiated from EBs with 5000 cells. Scale bar = 5 mm. D. Macroscopic views of cartilage tissue differentiated from EBs with 2000 cells. Scale bar = 5 mm. E. HE staining of cartilage tissues differentiated from EBs with 5000 cells. Scale bar = 500 µm. F. HE staining of cartilage tissues differentiated from EBs with 2000 cells. Scale bar = 500 µm. G. Alcian blue staining of cartilage tissues differentiated from EBs with 5000 cells. Scale bar = 500 µm. H. Alcian blue staining of cartilage tissues differentiated from EBs with 2000 cells. Scale bars = 500 µm.

First, to evaluate the effect of embryoid body (EB) size on chondrogenic differentiation, hESCs were seeded at 5000 or 2000 cells per well and cultured for 4 days. In floating culture, EBs composed of either 5000 or 2000 cells formed a single round sphere (Fig. 1B). The diameter of EBs composed of 5000 ESCs was 630 µm, while that of EBs composed of 2000 ESCs was 345 µm. The cartilage tissue from EBs consisting of 5000 cells was 7 mm in diameter, and that from EBs consisting of 2000 cells was 11 mm in diameter (Fig. 1A and 1D). The HE staining results showed that all chondrocytes were evenly distributed throughout the cartilage lumen (Fig. 1E and 1F). However, it was observed that cartilage tissues created from EBs consisting of 5000 cells contained other tissues such as epidermis, in addition to cartilage. In contrast, cartilage tissues accounted for a significantly higher percentage of tissues from EBs with 2000 cells (Fig. 1G and 1H). Therefore, the subsequent studies were conducted using EBs consisting of 2000 cells.

### 3.2. Low-density seeding of embryoid bodies (EBs) optimizes chondrogenic differentiation

To evaluate the impact of seeding density during EB adhesion culture on chondrogenic differentiation, we prepared cell sheets and cartilage tissue by seeding EBs at a density of 15.70 EB/cm^2^ or 12.21 EB/cm^2^ (Fig. 2A, 2B and 2C). The Alcian blue staining intensity percentage in cell sheets seeded with EBs at a high density (15.70 EB/cm^2^) was approximately 10% (Fig. 2D). In contrast, cell sheets seeded at a low density (12.21 EB/cm^2^) contained approximately 30% (Fig. 2D). The resulting cartilage tissue from high-density seeded EBs was 11 mm in size. In contrast, the cartilage tissue produced from the low-density seeded EBs was about 10 mm in size (Fig. 2E and 2F). Alcian blue staining revealed the presence of chondrocytes in both cartilage tissues (Fig. 2G and 2H).

**Figure 2.**
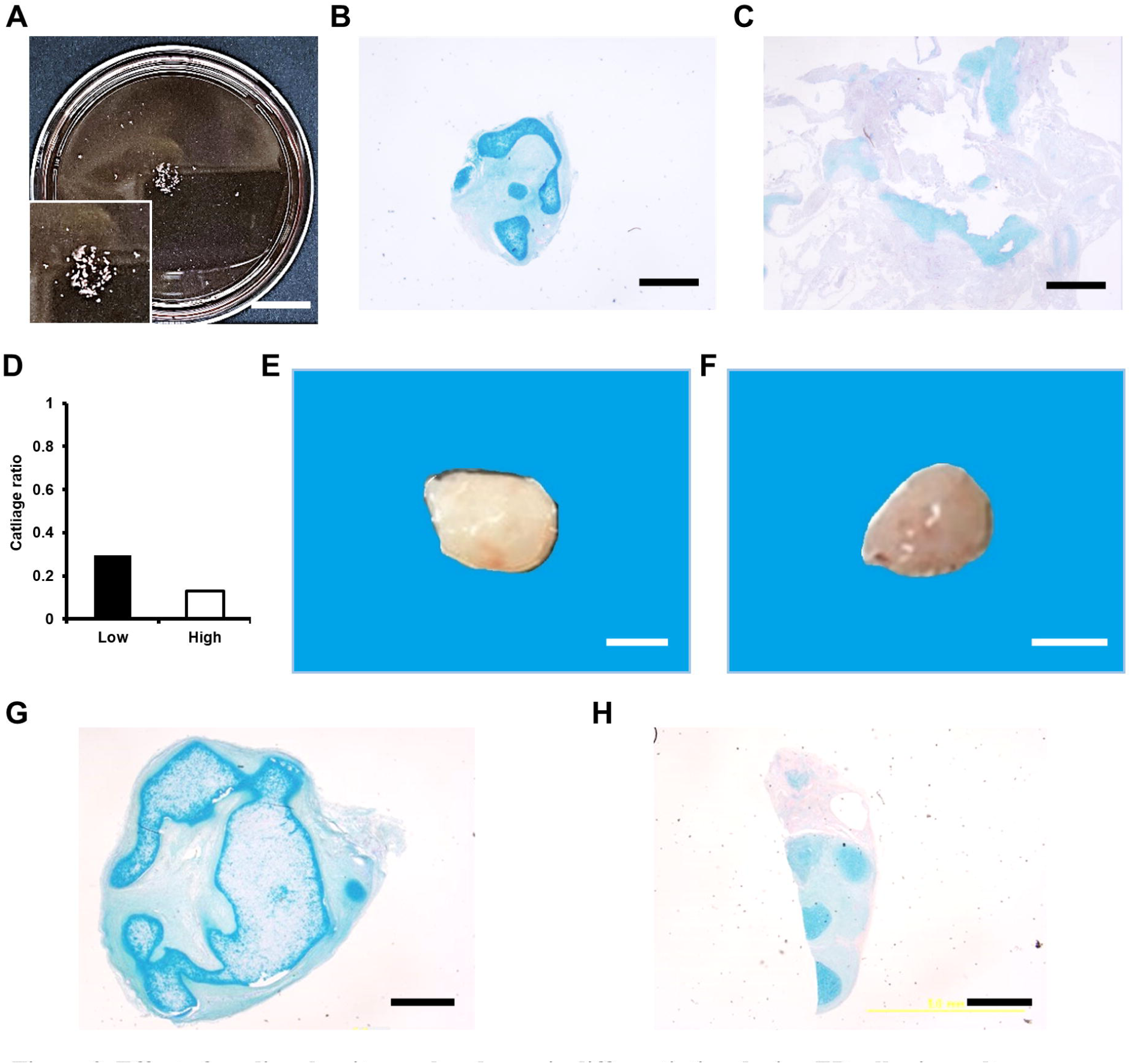
Effect of seeding density on chondrogenic differentiation during EB adhesion culture. A. Macroscopic views of culture dish after EB seeding. Inset: High-power view. Scale bar = 20 mm. B. Alcian blue staining of cell sheets at low EB seeding density (12.21 EB/cm^2^). Scale bar = 2 mm. C. Alcian blue staining of cell sheets at high EB seeding density (15.70 EB/cm^2^). Scale bar = 2 mm. D. Cartilage ratio of cell sheets at low EB seeding density (12.21 EB/cm^2^) or high EB seeding density (15.70 EB/cm^2^). E. Macroscopic views of cartilage tissue at low EB seeding density (12.21 EB/cm^2^). Scale bar = 5 mm. F. Macroscopic views of cartilage tissue at high EB seeding density (15.70 EB/cm^2^). Scale bar = 5 mm G. Alcian blue staining of cartilage tissue at low EB seeding density (12.21 EB/cm^2^). Scale bar = 2 mm. H. Alcian blue staining of cartilage tissue at high EB seeding density (15.70 EB/cm^2^). Scale bar = 2 mm.

### 3.3. Chondrogenic differentiation is promoted by the number of days in culture

To evaluate the effect of culture duration on chondrogenic differentiation, cell sheets were collected at 21, 30, or 60 days after the start of differentiation induction. The cell sheets prepared after 30 or 60 days of culture were transplanted into NOG mice to produce chondrocyte tissues. The cell sheets were not transplanted 21 days after the start of differentiation induction due to the absence of chondrocyte characteristics or staining with Alcian blue. The results showed that the cartilage tissue after 30 days of incubation was 3 mm, while the cartilage tissue after 60 days of incubation was 15 mm (Fig. 3A and 3B). Both tissues showed lacuna with chondrocytes, which is a characteristic of chondrocytes, but this was more pronounced in the 60-day culture (Fig. 3C and 3D). The 60-day culture period had a higher cell abundance (Fig. 3D). After the 60-day culture period, the cells were stained more intensely with Alcian blue (Fig. 3E and 3F). Gene expression analysis showed that the expression of cartilage markers, including ACAN, COL2A1, SOX9, and Vimentin, increased proportionally with the number of days of culture (Fig. 3G). Conversely, the gene expression of the undifferentiated markers OCT3/4 and NANOG tended to decrease with the duration of culture, indicating ongoing differentiation.

**Figure 3.**
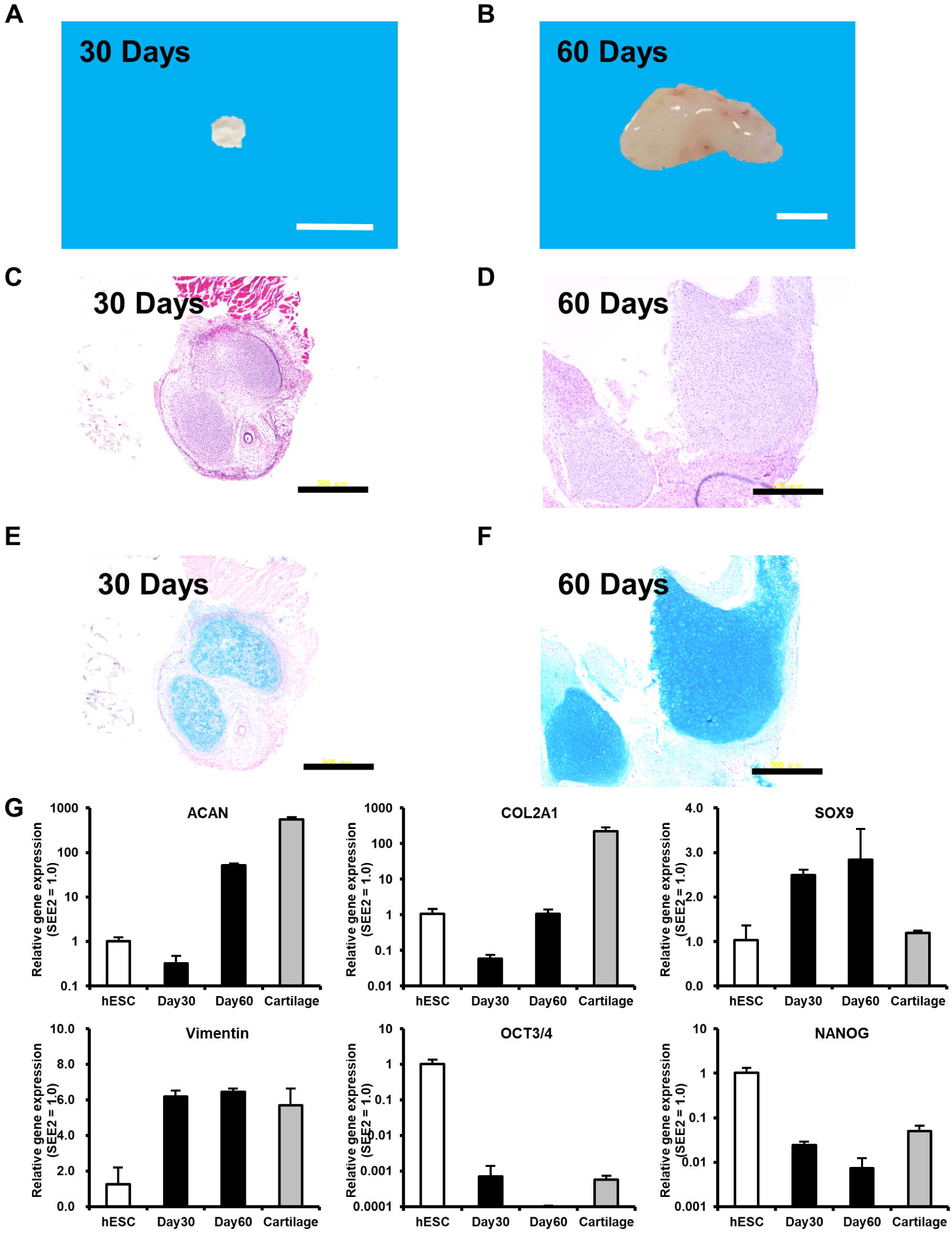
Effect number of days in culture on chondrogenic differentiation. A. Macroscopic views of cartilage tissue generated from cell sheets with 30 days of culture. Scale bar = 5 mm. B. Macroscopic views of cartilage tissue generated from cell sheets with 60 days of culture. Scale bar = 5 mm. C. HE staining cartilage tissue generated from cell sheets with 30 days of culture. Scale bar = 500 µm D. HE staining cartilage tissue generated from cell sheets with 60 days of culture. Scale bar = 500 µm. E. Alcian blue staining of cartilage tissue from cell sheets with 30 days of culture. Scale bars = 500 µm. F. Alcian blue staining of cartilage tissue from cell sheets with 60 days of culture. Scale bars = 500 µm. G. Gene expression analysis of cartilage tissues generated from cell sheets with different numbers of days in culture. The analysis was performed by qRT-PCR and the expression in ESCs was set as 1.0, and the relative values are shown. Data were normalized by the housekeeping gene UBC. Error bars indicated standard deviation.

In addition to in vitro culture conditions, in vivo transplantation conditions also have a significant impact on chondrogenic differentiation. The size of the cartilage tissue increased proportionally to the number of transplanted cell sheets. Specifically, the cartilage tissue produced by transplanting two cell sheets was 10 mm, three sheets was 13 mm, four sheets was 15 mm, and five sheets was 22 mm. The cartilage tissue reached its maximum maturity after 60 days of transplantation (Fig. S1A, S1B and S1C). The size of the cartilage tissue produced increased with in vivo time at 30, 60, and 90 days, and the intensity of Alcian blue staining reached its peak at 60 days (Fig. S2A, S2B and S2C). However, at 90 days, the cartilage tissue showed hypertrophic cartilage-like staining in Alcian blue staining, and differentiation into bone tissue and bone marrow was observed in the HE staining images.

### 3.4. Chondrogenic differentiation can be automated using a cell culture system

The process was transitioned from manual to automated using an automated cell culture system for competent and scalable chondrocyte production (Fig. 4A). Comparison of embryoid bodies (EBs) produced by either the automated cell culture system or the manual culture process showed that uniform EBs were produced with no difference in size or shape (Fig. 4B). The average recovery efficiency of EBs by the machine was 98% (94 EBs/96 wells-plate), which is a high level. The differentiation into cell sheets by adhesion culture was performed using an automated cell culture system (CellKeeper®). At 60 days after the start of differentiation induction, cartilage-like morphology was observed, similar to cell sheets produced by the manual culture process (Fig. 4C). No differences in size or shape were observed between the cartilage tissues prepared from either the automated cell culture system or the manual culture process (Fig. 4D and 4E). HE staining revealed lacuna with chondrocytes, a characteristic of chondrocytes in all cartilage tissues (Fig. 4F and 4G). Both cartilage tissues were stained with Alcian blue and showed similar staining intensity (Fig. 4H and 4I). The results indicate that the cartilage tissues produced by both the manual culture process and the automated cell culture system are composed of cartilage cells of comparable quality.

**Figure 4.**
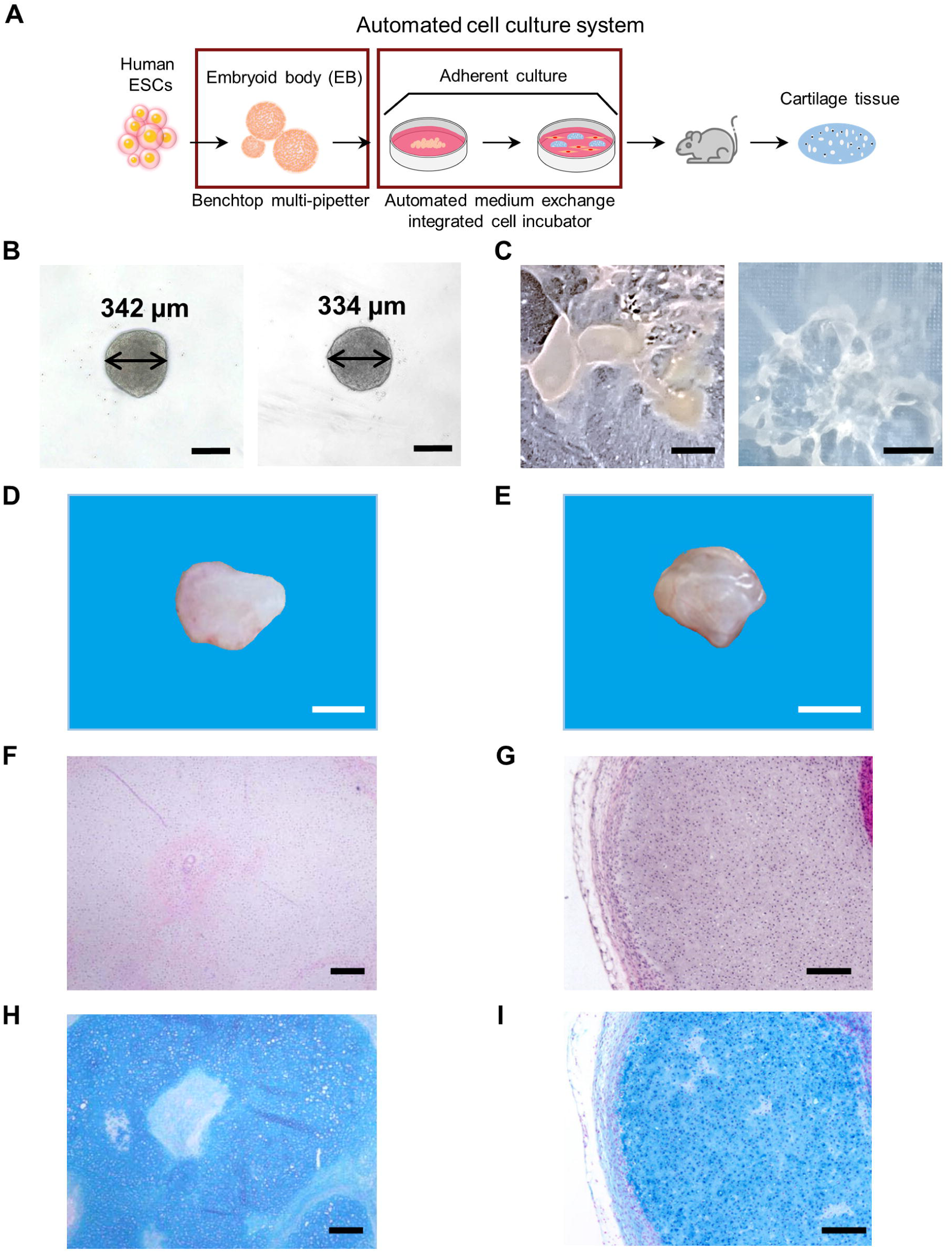
Automation of chondrogenic differentiation using a cell culture system. A. Schematic diagram of this study. Red boxes: automated sections. B. Phase contrast microscopy images of EBs prepared by an automated cell culture system (automated pipetting machine) or by manual cell culture process. Left: EBs prepared by manual cell culture process; right: EBs prepared by automated pipetting machine; Scale bars = 200 µm. C. Macroscopic views of cell sheets produced by an automated cell culture system or by manual cell culture process. left: cell sheet produced by manual cell culture process. right: cell sheet produced by an automated pipetting machine. Right: Cell sheet produced by an automated cell culture system, Scale bars = 5 mm. D. Macroscopic views of cartilage tissue prepared from cell sheets produced by manual cell culture process-generated cell sheets. Scale bar = 5 mm. E. Macroscopic views of cartilage tissue prepared from cell sheets produced by a cell sheet produced by an automated cell culture system. Scale bar = 5 mm. F. HE staining of cartilage tissue prepared from cell sheets produced by manual cell culture process-generated cell sheet. Scale bar = 200 µm. G. HE staining of cartilage tissue prepared from cell sheets produced by a cell sheet produced by an automated cell culture system. Scale bar = 200 µm. H. Alcian blue staining of cartilage tissue prepared from cell sheets produced by manual cell culture process-generated cell sheet. Scale bar = 200 µm. I. Alcian blue staining of cartilage tissue prepared from cell sheets produced by a cell sheet produced by an automated cell culture system. Scale bar = 200 µm.

### 3.5. Dome-shaped cartilage formation in vitro

Analysis of Alcian blue staining revealed that the cell sheet contained cartilage and non-cartilage areas (Fig. 5A and 5B). Cartilage was randomly generated on the cell sheet as “islands.” The non-cartilage areas consisted of one or about two layers of fibroblasts, with no ECM secretion (Fig. 5C and S3A). The cartilage areas had a well-developed ECM, lacuna with chondrocytes, and several layers of chondrocytes (Fig. 5C, 5D and S3B). Histological analysis showed that the cartilage had a half-moon shape on the dish and several layers of stromal-like cells surrounding the cartilage. On the other hand, no cells other than chondrocytes were found inside the cartilage (Fig. 5D). It appeared that chondroblasts were in the layer of stromal-like cells surrounding the cartilage.

**Figure 5.**
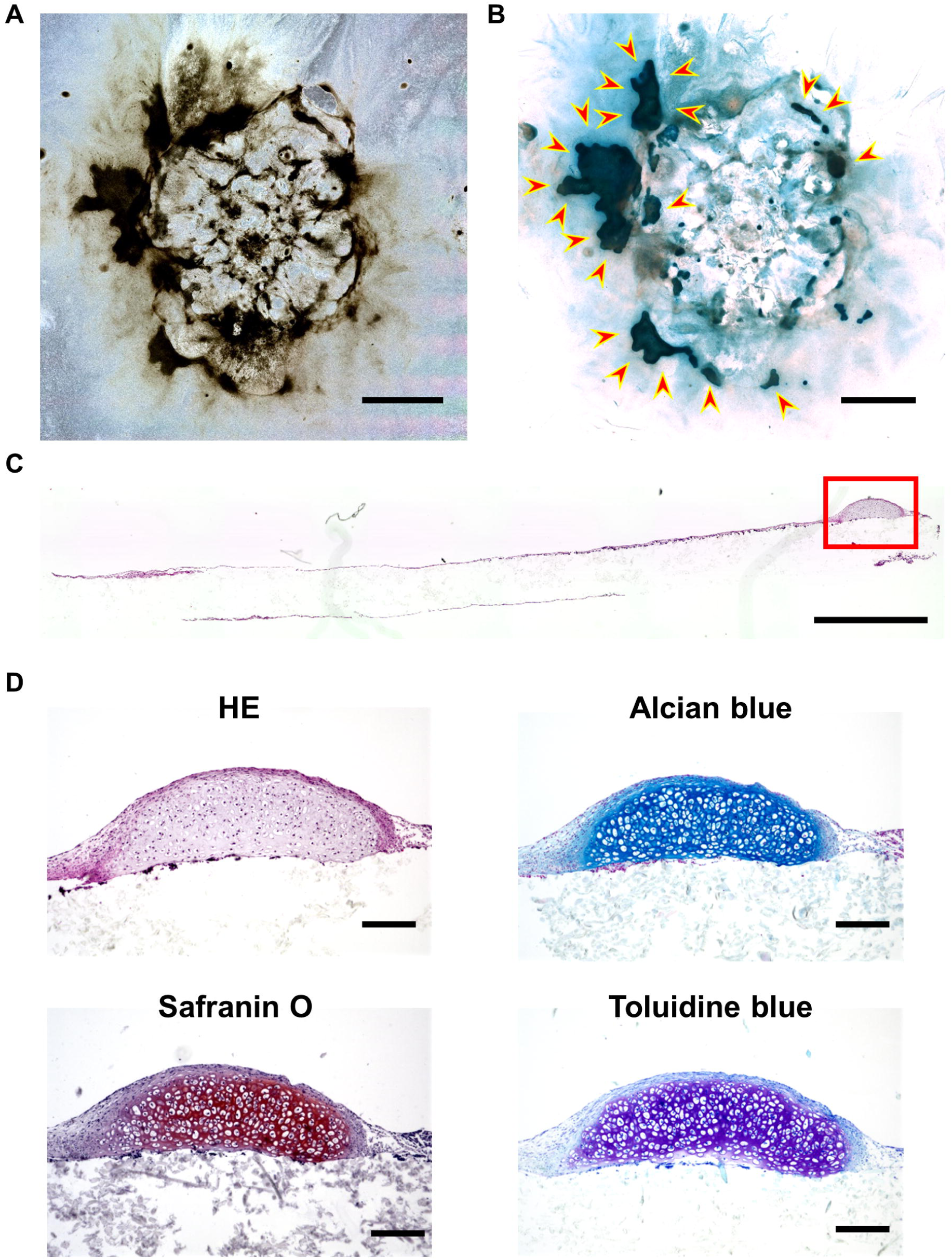
Characteristics of histology and location of cartilage in cell sheets. A. Phase contrast microscopy images of a cell sheet for culture 60 days. Scale bar = 10 mm. B. Bright-field image of Fig. 5A after alcian blue staining. Arrows: cartilage, areas darkened by alcian blue. Scale bar = 10 mm. C. HE staining of a vertical section of a cell sheet for culture 60 days. Red frame: cartilage; Scale bar = 1 mm. D. Various histological stains of the red box in Fig. 5C. Top left: HE staining, top right: Alcian blue staining, bottom left: Safranin O staining, bottom right: Toluidine blue staining. Scale bars = 200 µm.

### 3.6. Translucency-based assessment of cartilage quality

Because cartilage has a low number of cells per unit volume, we were able to distinguish cartilage on the cell sheet by the difference in light transmission under the stereomicroscope (Fig. 6A). In addition, because cartilage has physical strength and elasticity, only cartilage could be harvested from the cell sheet (Fig. 6B). The entire cell sheet was transplanted into NOG mice, and cartilage tissue was obtained (Fig. 6C). Histological analysis revealed that in addition to chondrocytes, intestinal epithelial-like cells, cardiomyocyte-like cells, and skin-like cells were also found in the cartilage tissue. Some cartilage tissue also underwent endochondral ossification (Fig. 6E). Only the cartilage component of the cell sheets was transplanted into NOG mice, and cartilage tissue was obtained (Fig. 6D). Histological analysis showed that the cartilage tissue consisted of uniform chondrocytes (Fig. 6F). These results suggest that cartilage ossification was caused by other cells included in the transplantation.

**Figure 6.**
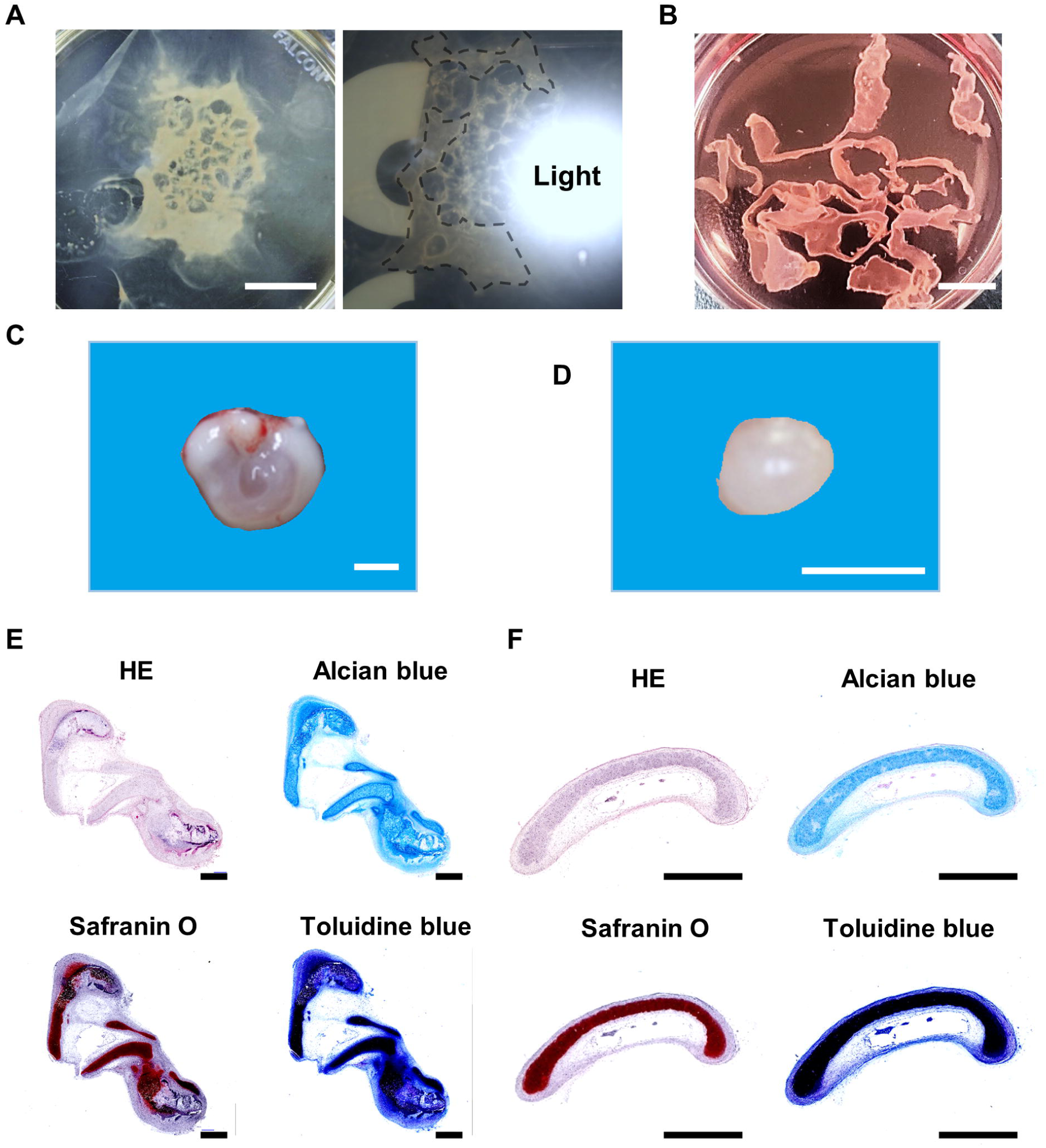
Effect of cartilage purity on cartilage tissue production. A. Macroscopic views of a cell sheet for culture 60 days. Left: normal light, Right: light from behind. Black dotted line: cartilage. Scale bar = 20 mm. B. Macroscopic views of cartilage harvested from cell sheets. Scale bar = 10 mm. C. Macroscopic views of cartilage tissue produced from a whole cell sheet. Scale bar = 5 mm. D. Macroscopic views of cartilage tissue produced from cartilage harvested from cell sheets. Scale bar = 5 mm. E. Various histological stains of Fig. 6C. Top left: HE staining, top right: Alcian blue staining, bottom left: Safranin O staining, bottom right: Toluidine blue staining. Scale bars = 10 mm. F. Various histological stains of Fig. 6D. Top left: HE staining, top right: Alcian blue staining, bottom left: Safranin O staining, bottom right: Toluidine blue staining. Scale bars = 10 mm.

## 4. Discussion

This study established a protocol for obtaining cartilage tissue derived from human embryonic stem cells (hESCs) and identified the culture conditions that affect cartilage differentiation. This culture system provides a simple method for obtaining cartilage tissue for regenerative medicine and a valuable model for studying the conditions for chondrogenic differentiation.

EBs are a useful in vitro model for differentiation. However, optimizing the number of cells in EBs is essential to differentiate target cells efficiently. Several studies have addressed the correlation between EB size and cell differentiation fate[21–26] and ESCs do not differentiate into cardiomyocytes when the number of constituent cells in the EB is less than 1000[24]. The cell sheets prepared in this study with 2000 EBs contain cardiomyocyte-like cells with the same mesodermal origin as cartilage, and the cell sheets rarely contract. Setting the number of constituent cells of EBs to 500-1000 may make cartilage differentiation more efficient. However, this study also revealed that the seeding density of EBs affects chondrogenic differentiation. Cartilage formation was observed in the cell sheet when EBs were seeded clustered at the center of the dish. On the other hand, seeding EBs evenly did not result in cartilage formation, indicating a correlation between EB seeding density and chondrogenic differentiation factors for MSCs, cell density, and tension between cells. Chondrogenic differentiation of mesenchymal stem cells (MSCs) requires a high cell density, as exemplified by pellet culture[27]. Additionally, MSCs reported to differentiate into chondrocytes under muscular tension[28]. The cartilage of the cell sheets did not form at the point of EB attachment but instead formed in a circle approximately 10 mm away from the attachment point. These findings suggest that MSCs within this region had an optimal cell density and differentiated into chondrocytes due to tension stress. In summary, when considering a smaller number of EB constructs (500-1000 cells), it may be necessary to increase the seeding density due to reduced cell density and tension.

The size of cartilage tissue can be ensured by mass culturing cell sheets using an automated cell culture system. The irregular shape of cartilage tissue is a problem that needs to be solved to avoid inefficient use of cartilage tissue. It is possible to shape cartilage tissue by surrounding it with materials such as mesh or plastic to limit the range of cartilage growth. This study is the first to report optical features specific to cartilage. By combining this knowledge with image recognition artificial intelligence (AI), the process of sorting cartilage from cell sheets may automated. The ability to harvest only cartilage from the cell sheet can produce uniform cartilage tissue, ensuring strength and reducing the possibility of teratoma formation. PluriSIn can reduce the risk of teratoma formation by removing undifferentiated pluripotent stem cells[29,30].

## 5. Conclusion

This study shows that human ESCs generated cartilage in adhesion culture. In addition, we revealed the optimal cell numbers of EB, differentiation period, and cell density in chondrogenic differentiation. Combining these findings with an automated cell culture system shows the potential to produce cartilage of a size that can be used in human clinical practice. This cartilage tissue has potential applications used clinically to repair organs such as the trachea, joints, ears, and nose.

## Declarations

### Ethics approval and consent to participate

All experiments involving handling of human cells and tissues were approved by the Institutional Review Board at the National Center for Child Health and Development (2021-178). In compliance with the Declaration of Helsinki, informed consent was obtained from all tissues and cell donors. When the donors were under 18, informed consent was obtained from parents. Human cells in this study were utilized in full compliance with the Ethical Guidelines for Medical and Health Research Involving Human Subjects (Ministry of Health, Labor, and Welfare (MHLW), Japan; Ministry of Education, Culture, Sports, Science and Technology (MEXT, Japan) and performed in full compliance with the Ethical Guidelines for Clinical Studies (Ministry of Health, Labor, and Welfare, Japan). The cells were banked after approval of the Institutional Review Board at the National Institute of Biomedical Innovation (May 9, 2006). The derivation and cultivation of human embryonic stem cell (hESC) lines were performed in full compliance with “the Guidelines for Derivation and Distribution of Human Embryonic Stem Cells (Notification of MEXT, No. 156 of August 21, 2009; Notification of MEXT, No. 86 of May 20, 2010) and “the Guidelines for Utilization of Human Embryonic Stem Cells (Notification of MEXT, No. 157 of August 21, 2009; Notification of MEXT, No. 87 of May 20, 2010)”. All procedures of animal experiments were approved by the Institutional Animal Care and Use Committee in National Center for Child Health and Development, based on the basic guidelines for the conduct of animal experiments in implementing agencies under the jurisdiction of the Ministry of Health, Labour and Welfare (Notification of MHLW, No. 0220-1 of February 20, 2015). The Institutional Animal Care and Use Committee of the National Center for Child Health and Development approved the protocols of the animal experiments (approval number: A2003-002-C20-M01, title: Research on regenerative medicine and cell medicine using mesenchymal stem cells, iPS cells, and ESCs, and on toxicity testing). Animal experiments were performed according to protocols approved by the Institutional Animal Care and Use Committee of the National Research Institute for Child Health and Development.

### Consent for publication

Not applicable.

### Availability of data and material

The datasets and cells used during the current study are available from the corresponding author on reasonable request.

### Authors’ contributions

AU, YO, AT, OK, KT and CJ designed the experiments. OK, KT, AT, RI, CJ, HY, HK and YO performed the experiments. OK, KT, AT, CJ, SA, YO, HY and MK analyzed data. MK, HK, YF, SE, HA and AU contributed to the reagents, tissues and analysis tools. OK, CJ, SA, MI, SE, HA and AU discussed the data and manuscript. CJ, SA and AU wrote this manuscript. All authors read and approved the final manuscript.

## Acknowledgments

We would like to express our sincere thanks to K. Miyado and M. Komura for fruitful discussion, to M. Ichinose for providing expert technical assistance, to C. Ketcham for English editing and proofreading, and to E. Suzuki and K. Saito for secretarial work.

