## Supplementary figures and images for "Automated Xeno-Free Chondrogenic Differentiation from Human Embryonic Stem Cells: Enhancing Efficiency and Ensuring High-Quality Mass Production"

### Fig. S1

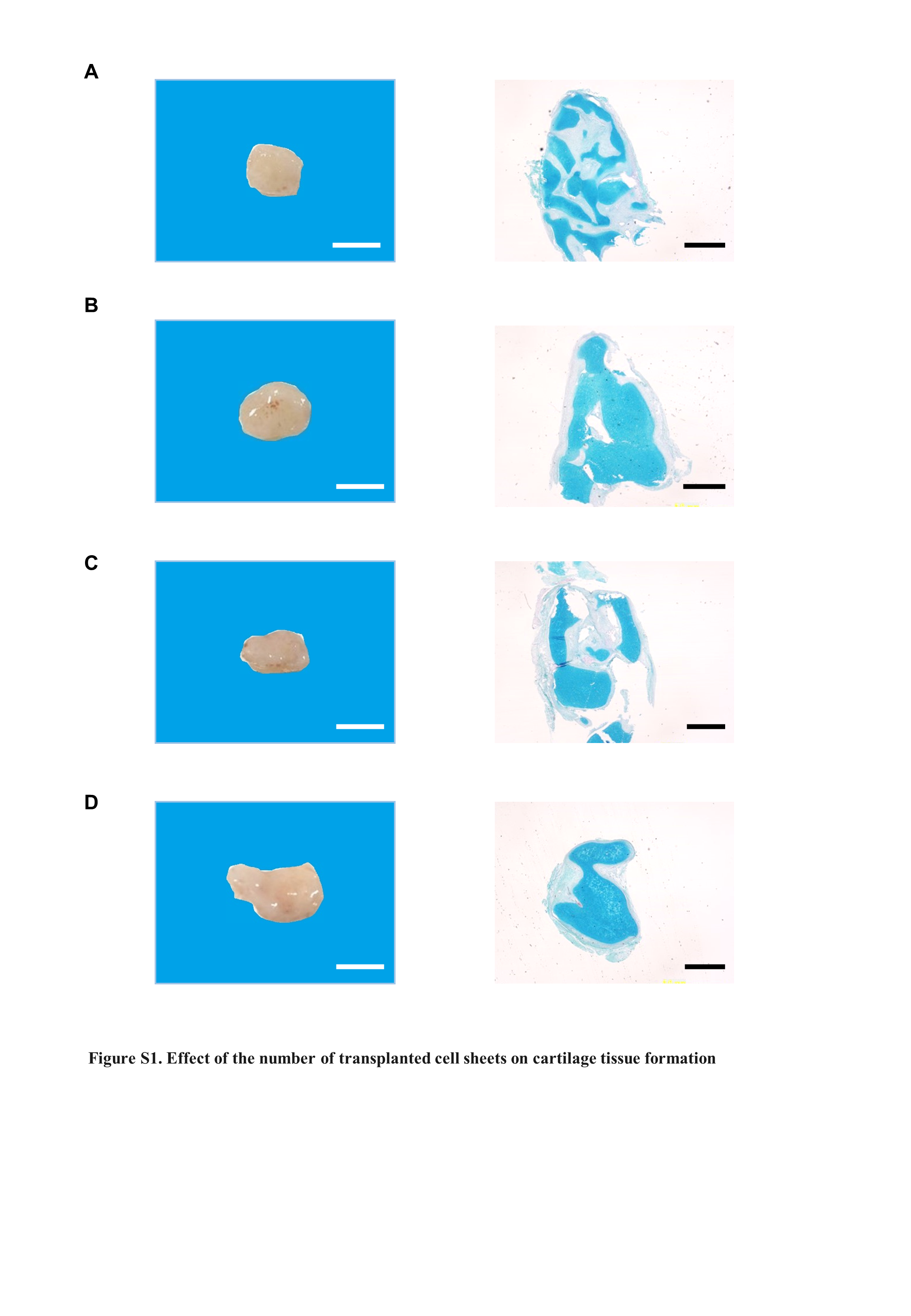

### Fig. S2

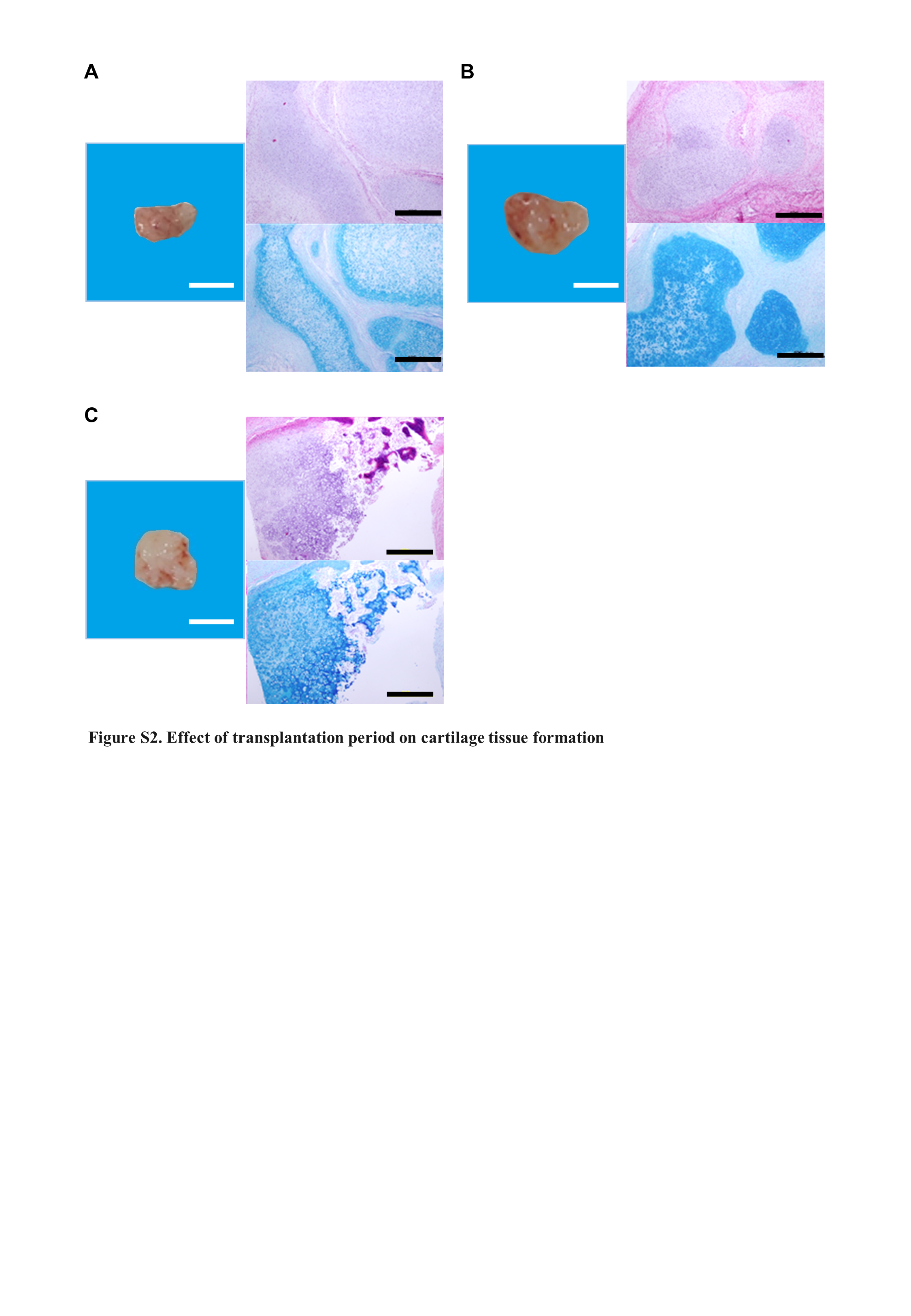

### Fig. S3

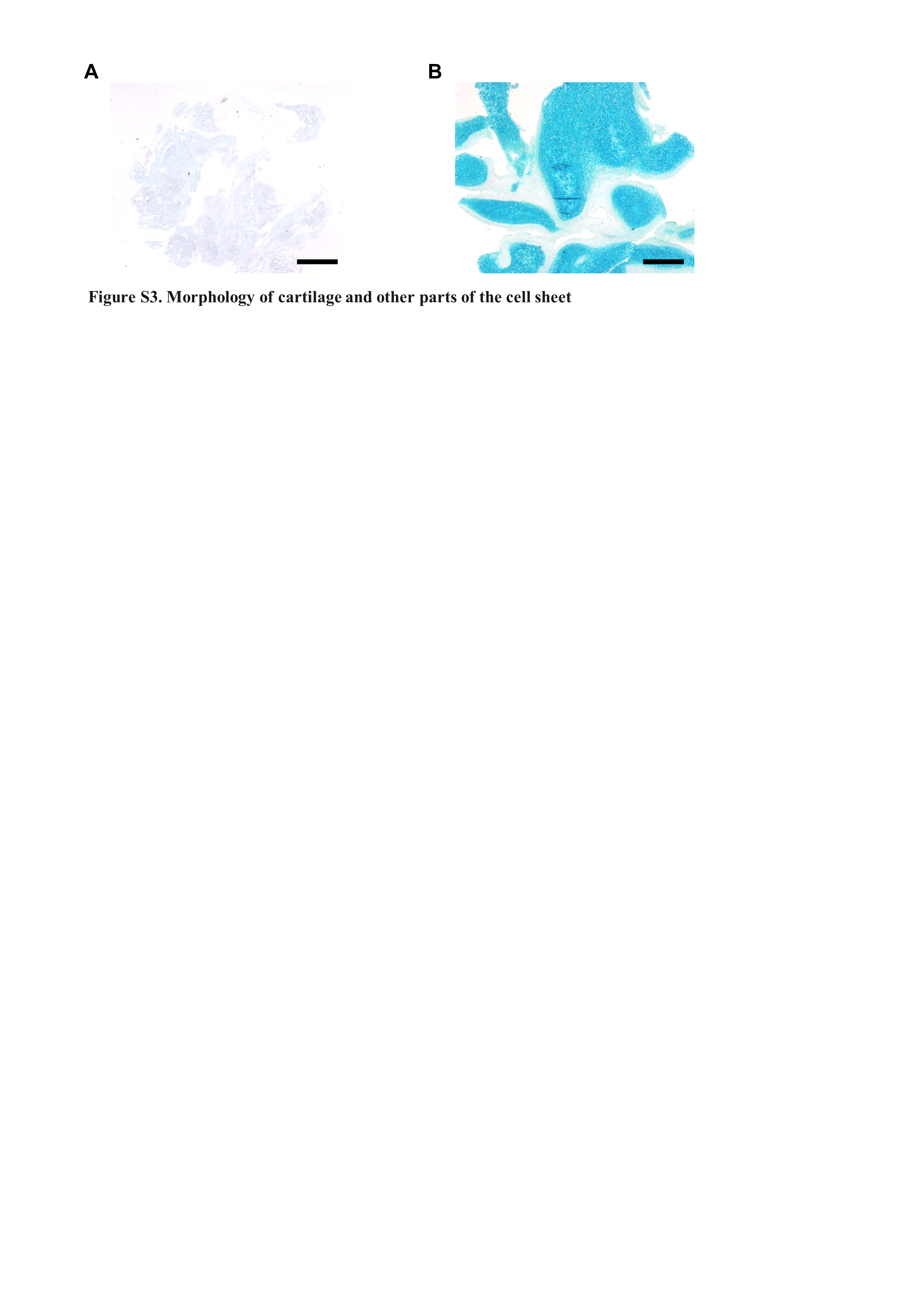
